# Variant SARS-CoV-2 mRNA vaccines confer broad neutralization as primary or booster series in mice

**DOI:** 10.1101/2021.04.13.439482

**Authors:** Kai Wu, Angela Choi, Matthew Koch, Sayda Elbashir, LingZhi Ma, Diana Lee, Angela Woods, Carole Henry, Charis Palandjian, Anna Hill, Hardik Jani, Julian Quinones, Naveen Nunna, Sarah O’Connell, Adrian B McDermott, Samantha Falcone, Elisabeth Narayanan, Tonya Colpitts, Hamilton Bennett, Kizzmekia S Corbett, Robert Seder, Barney S Graham, Guillaume BE Stewart-Jones, Andrea Carfi, Darin K Edwards

## Abstract

Severe acute respiratory syndrome coronavirus 2 (SARS-CoV-2) is the causative agent of a global pandemic. Safe and effective COVID-19 vaccines are now available, including mRNA-1273, which has shown 94% efficacy in prevention of symptomatic COVID-19 disease. However, the emergence of SARS-CoV-2 variants has led to concerns of viral escape from vaccine-induced immunity. Several variants have shown decreased susceptibility to neutralization by vaccine-induced immunity, most notably B.1.351 (Beta), although the overall impact on vaccine efficacy remains to be determined. Here, we present the initial evaluation in mice of 2 updated mRNA vaccines designed to target SARS-CoV-2 variants: (1) monovalent mRNA-1273.351 encodes for the spike protein found in B.1.351 and (2) mRNA-1273.211 comprising a 1:1 mix of mRNA-1273 and mRNA-1273.351. Both vaccines were evaluated as a 2-dose primary series in mice; mRNA-1273.351 was also evaluated as a booster dose in animals previously vaccinated with mRNA-1273. The results demonstrated that a primary vaccination series of mRNA-1273.351 was effective at increasing neutralizing antibody titers against B.1.351, while mRNA-1273.211 was effective at providing broad cross-variant neutralization. A third (booster) dose of mRNA-1273.351 significantly increased both wild-type and B.1.351-specific neutralization titers. Both mRNA-1273.351 and mRNA-1273.211 are being evaluated in pre-clinical challenge and clinical studies.

## 1. Introduction

Since the declaration of a global pandemic by the World Health Organization on March 11, 2020, infection with the severe acute respiratory syndrome coronavirus 2 (SARS-CoV-2) has led to approximately 4.7 million deaths worldwide [1, 2]. Shortly after the SARS-CoV-2 genetic sequence was determined, mRNA-1273—a novel lipid nanoparticle (LNP) encapsulated messenger RNA-based vaccine encoding for a prefusion-stabilized, full-length spike (S) glycoprotein of the Wuhan-Hu-1 isolate of SARS-CoV-2—was developed [3, 4]. Vaccination with two 100 μg doses of mRNA-1273 four weeks apart was 94% efficacious against symptomatic COVID-19 disease; mRNA-1273 was granted Emergency Use Authorization by the Food and Drug Administration in December 2020 [5, 6].

The emergence of SARS-CoV-2 variants with substitutions in the receptor binding domain (RBD) and N-terminal domain (NTD) of the viral S protein has raised concerns among scientists and health officials [7–10]. The entry of coronaviruses into host cells is mediated by interaction between the RBD of the viral S protein and the host receptor, angiotensin-converting enzyme 2 (ACE2) [3, 11–14]. Several studies have shown that the RBD is the main target of neutralizing antibodies against SARS-CoV-2 [4, 15–17]. A neutralization “supersite” has also been identified in the NTD [18]. A decrease in vaccine-mediated viral neutralization has been correlated with amino acid substitutions in the RBD (eg, K417T/N, E484K, and N501Y) and NTD (eg, L18F, D80A, D215G, and Δ242-244) of the S protein. Some of the most recently circulating variants of concern (VOCs) and variants of interest (VOIs) with key mutations in the RBD and NTD— including B.1.1.7 (Alpha), B.1.351 (Beta), P.1 (Gamma), B.1.526 (Iota), and B.1.427/B.1.429 (Epsilon or CAL.20C) lineages—have shown reduced susceptibility to neutralization from convalescent serum and resistance to monoclonal antibodies [18–25]. Note that mutations in the NTD domain, specifically the neutralization supersite, are extensive in the B.1.351 lineage virus [18].

Using 2 orthogonal pseudovirus neutralization (PsVN) assays based on vesicular stomatitis virus (VSV) and lentivirus expressing S variants, neutralizing capacity of sera from phase 1 participants and non-human primates (NHPs) that received 2 doses of mRNA-1273 was reported [26]. No significant impact on neutralization against the B.1.1.7 variant was observed. However, reduced neutralization was measured against the B.1.351 variant, and to a lesser extent, in the P.1 B.1.427/B.1.429 and B.1.1.7+E484K variants [26]. Clinical studies in South Africa demonstrated reduced efficacy against symptomatic COVID-19 disease for the NVX-CoV2373 (Novavax), AZD1222 (University of Oxford/AstraZeneca), and Ad26.COV2.S (Janssen/Johnson & Johnson) vaccines [27–30]. Pfizer/BioNTech has recently reported high efficacy of the mRNA BNT162b2 vaccine against B.1.351 among a small number of recipients from South Africa in the phase 2–3 portion of a global phase 1–2–3 trial; however, a report from Israel suggests increased breakthrough infection rates by B.1.351 in BNT162b2 vaccinated individuals at 7–14 days after the second dose [31, 32]. Studies have demonstrated reduced neutralization titers against the full B.1.351 variant following mRNA-1273 vaccination, although levels are still significant and expected to be protective based on challenge studies in NHPs [26, 33, 34]. Despite this prediction of continued efficacy of mRNA-1273 against this key variant of concern (VOC), the magnitude and duration of vaccine-mediated protection is still unknown. Moreover, a key related question is whether development of new mRNA vaccines to match the B.1.351 variant will enable enhanced neutralization responses and durability.

Herein, we present the design and pre-clinical evaluation of updated mRNA-1273 vaccines against SARS-CoV-2 variants, which include monovalent mRNA-1273.351 and a multivalent mRNA-1273.211. Like mRNA-1273, mRNA-1273.351 encodes the prefusion stabilized S protein of SARS-CoV-2; however, in contrast to mRNA-1273, mRNA-1273.351 incorporates key mutations present in the B.1.351 variant, including L18F, D80A, D215G, Δ242-244, R246I, K417N, E484K, N501Y, D614G, and A701V (Fig. 1). To expand the breadth of coverage to multiple circulating variants as well as the ancestral wild-type virus that is still circulating globally, mRNA-1273.211 is a 1:1 mix of mRNA-1273 and mRNA-1273.351. Two initial studies were performed in BALB/c mice to evaluate the immunogenicity of mRNA-1273.351 or mRNA-1273.211 as a primary series and/or as a booster. The first study assessed the immunogenicity of a primary series (day 0, 21) of mRNA-1273.351 or mRNA-1273.211 versus mRNA-1273. A second study evaluated the immunogenicity of a third (booster) dose of mRNA-1273.351 213 days after mice were vaccinated with a primary series of mRNA-1273, and immunogenicity was assessed before and after the booster dose.

**Fig. 1.**
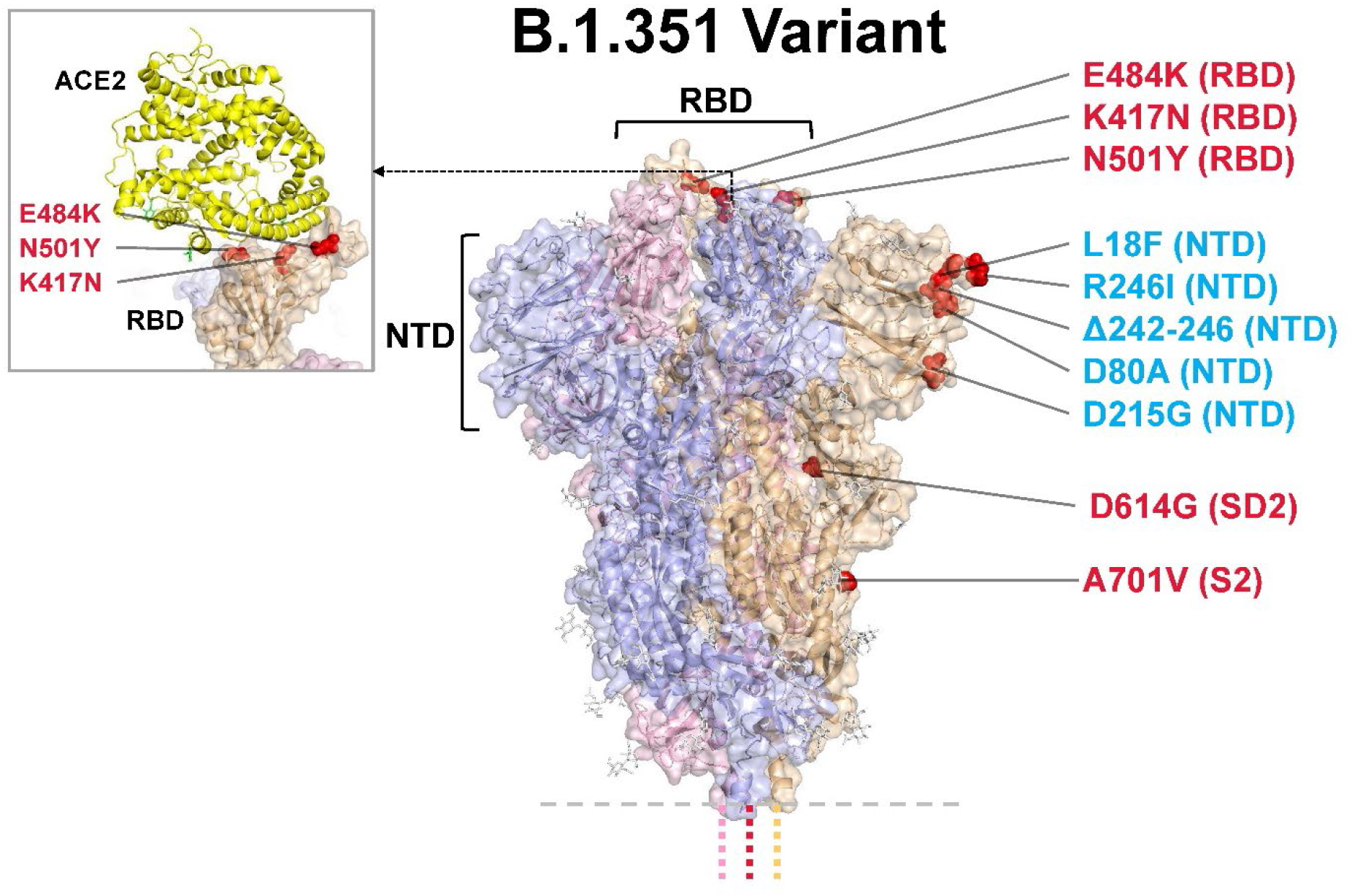
Model of S protein. mRNA-1273.351 encodes the B.1.351 lineage S variant. Surface representation of the trimeric S protein in the vertical view with the locations of surface-exposed mutated residues highlighted in red spheres and labelled on the gray monomer. The inset shows superimposition of ACE2 receptor domain and the RBD. S protein structure, 6VSB [3]. ACE2-RBD structure, 6M0J [44]. ACE2 = angiotensin converting enzyme 2; NTD = N-terminal domain; RBD = receptor binding domain.

## 2. Methods

### 2.1 Data reporting

No statistical methods were used to predetermine sample size. The experiments were not randomized and the investigators were not blinded to allocation during experiments and outcome assessment.

### 2.2 Pre-clinical vaccine mRNA and LNP production process

A sequence-optimized mRNA encoding prefusion-stabilized Wuhan-Hu-1 or B.1.351-variant SARS-CoV-2 S-2P protein was synthesized in vitro using an optimized T7 RNA polymerase-mediated transcription reaction with complete replacement of uridine by N1m-pseudouridine [35]. The reaction included a DNA template containing the immunogen open-reading frame flanked by 5’ untranslated region (UTR) and 3’ UTR sequences and was terminated by an encoded polyA tail. After transcription, the cap-1 structure was added to the 5’ end using the *Vaccinia* capping enzyme (New England Biolabs) and *Vaccinia* 2’-O-methyltransferase (New England Biolabs). The mRNA was purified by oligo-dT affinity purification, buffer exchanged by tangential flow filtration into sodium acetate, pH 5.0, sterile filtered, and kept frozen at −20°C until further use.

The mRNA was encapsulated in an LNP through a modified ethanol-drop nanoprecipitation process described previously [36]. Ionizable, structural, helper, and polyethylene glycol lipids were briefly mixed with mRNA in an acetate buffer, pH 5.0, at a ratio of 2.5:1 (lipids:mRNA). The mixture was neutralized with Tris-HCl, pH 7.5, sucrose was added as a cryoprotectant, and the final solution was sterile-filtered. Vials were filled with formulated LNP and stored frozen at −70°C until further use. The drug product underwent analytical characterization—which included the determination of particle size and polydispersity, encapsulation, mRNA purity, doublestranded RNA content, osmolality, pH, endotoxin, and bioburden—and the material was deemed acceptable for in vivo study.

### 2.3 Mouse model

Animal experiments were carried out in compliance with approval from the Animal Care and Use Committee of Moderna Inc. Female BALB/c mice (6 to 8 weeks old; Charles River Laboratories) were used. mRNA formulations were diluted in 50 μL of 1X phosphate-buffered saline (PBS), and mice were inoculated via intramuscular injection into the same hind leg for both prime, boost, and third dose. Control mice received PBS because prior studies have demonstrated that tested mRNA formulations do not create significant levels of non-specific immunity beyond a few days [37–39]. Sample size for animal experiments was determined on the basis of criteria set by the institutional Animal Care and Use Committee. Experiments were neither randomized nor blinded.

### 2.4 Enzyme-linked Immunosorbent Assay (ELISA)

Microtiter plates (96-well; Thermo) were coated with 1 μg/mL S-2P protein (Genscript) corresponding to the S protein of the Wuhan-Hu-1 virus. After overnight incubation at 4°C, plates were washed four times with PBS/0.05% Tween-20 and blocked for 1.5 hours at 37°C (SuperBlock-Thermo). After washing, five-fold serial dilutions of mouse serum were added (assay diluent: 0.05% Tween-20 and 5% goat serum in PBS). Plates were incubated for 2 hours at 37°C, washed and horseradish peroxidase-conjugated goat anti-mouse immunoglobulin G (IgG) (Southern Biotech) was added at a 1:20,000 dilution (S-2P) in assay diluent. Plates were incubated for 1 hour at 37°C, washed, and bound antibody was detected with a 3,3’,5,5’-tetramethylbenzidine (TMB) substrate (SeraCare). After incubation for 10 minutes at room temperature, the reaction was stopped by adding a TMB stop solution (SeraCare) and the absorbance was measured at 450 nm. Titers were determined using a four-parameter logistic curve fit in Prism v.8 (GraphPad Software, Inc.) and defined as the reciprocal dilution at approximately optical density 450 = 1.0 (normalized to a mouse standard on each plate).

### 2.5 Recombinant VSV-based PsVN assay

Codon-optimized full-length S protein of the original Wuhan-Hu-1 variant with D614G mutation (D614G) or the indicated S variants, listed in Table 1, were cloned into a pCAGGS vector. To make SARS-CoV-2 full-length S pseudotyped recombinant VSV-ΔG-firefly luciferase virus, BHK-21/WI-2 cells (Kerafast) were transfected with the S expression plasmid and subsequently infected with VSVΔG-firefly-luciferase as previously described [40]. For a neutralization assay, serially diluted serum samples were mixed with pseudovirus and incubated at 37°C for 45 minutes. The virus-serum mix was subsequently used to infect A549-hACE2-TMPRSS2 cells [41] for 18 hours at 37°C before adding ONE-Glo reagent (Promega) for measurement of the luciferase signal by relative luminescence units (RLUs). The percentage of neutralization was calculated based on the RLUs of the virus-only control, and subsequently analyzed using four-parameter logistic curve (Prism v.8).

**Table 1.** S-protein substitutions in SARS-CoV-2 variants evaluated in this study.

| Variant Name | Amino Acid Changes in S Protein Relative to Wuhan-Hu-1 |
| --- | --- |
| D614G | D614G |
| B.1.351 (Beta)<br>(501Y.V2) | L18F, D80A, D215G, Δ242-244, R246I, K417N, E484K,<br>N501Y, D614G, A701V |
| P.1 (Gamma)<br>(501Y.V3) | L18F, T20N, P26S, D138Y, R190S, K417T, E484K, N501Y,<br>D614G, H655Y, T1027I, V1176F |
| B.1.427/B.1.429 (Epsilon)<br>(452R.V1, CAL.20C) | S13I, W152C, L452R, D614G |

### 2.6 Statistical Analysis

Animal studies were completed once with all in vitro testing completed in duplicate or triplicate with 1 replicate, unless otherwise stated. Two-sided Wilcoxon matched-pairs signed rank test was used to compare the same animals against different viruses or at different time points. Statistical analyses were performed (Prism v.8). Geometric mean titers with 95% CIs and lower limits of detection are included, where applicable.

## 3. Results

### 3.1 mRNA-1273.351 and mRNA-1273.211 were immunogenic as a primary vaccination series in mice

The immunogenicity of mRNA-1273.351 and mRNA-1273.211 vaccines against the original Wuhan-Hu-1 variant with the D614G mutation (referred to as wild-type) and the B.1.351 variant was evaluated in BALB/c mice 2 weeks after the first and second injection (Fig. 1). Animals were vaccinated with 1 or 10 μg of mRNA-1273, mRNA-1273.351, or mRNA-1273.211 on a 0, 21-day schedule (Fig. 2a). S protein-binding antibody titers were assessed using a Wuhan-Hu-1 S-2P ELISA. In addition, neutralizing antibody titers were assessed in a VSV-based PsVN assay against wild-type and variant viruses (Table 1).

**Fig. 2.**
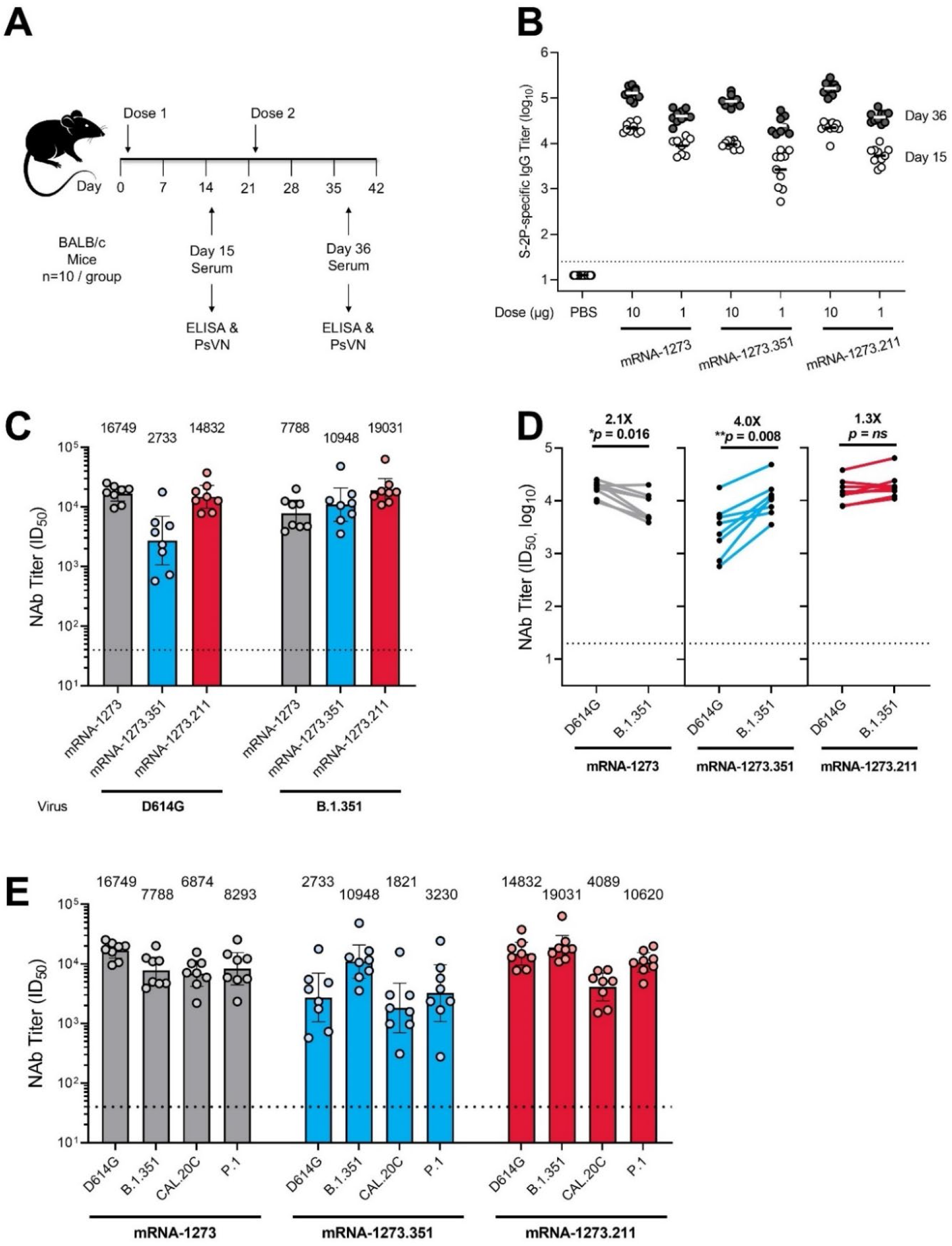
S protein-binding antibody and neutralization of variant SARS-CoV-2 pseudoviruses by serum from vaccinated BALB/c mice. **a,** BALB/c mice were immunized on a two-dose schedule with 1 or 10 μg mRNA-1273, mRNA-1273.351, mRNA-1273.211 (1:1 mix of mRNA-1273 and mRNA-1273.351), or PBS. **b,** Results from individual mouse sera (n = 10) following dose 1 (day 15) and after dose 2 (day 36) are represented as dots on each figure; the horizontal line indicates the GMT. **c,** BALB/c mice were immunized with 1 μg mRNA-1273, mRNA-1273.351, mRNA-1273.211. Each bar indicates the GMT value after dose 2 (day 36), which is listed as text above each plot. **d**, GMT values from individual mouse sera (n = 8 per antigen [randomly selected]) after dose 2 (day 36) of 1 μg mRNA-1273, mRNA-1273.351, or mRNA-1273.211 are represented as dots on each figure, with lines connecting matched pairs for the D614G and B.1.351 neutralization titers. Fold difference in neutralization against each virus was shown in text above each plot. Wilcoxon matched-pairs signed-rank test. Two-tailed p values: *<0.1; **<0.01. **e,** BALB/c mice were immunized with 1 μg mRNA-1273, mRNA-1273.351, or the multivalent mRNA-1273.211 vaccine (n = 8). Each bar indicates the GMT value after dose 2 (day 36), which is listed as text above each plot. The horizontal dotted line indicates the lower limit of quantitation for NAb titer at 40 ID_50_. ELISA = enzyme-linked immunosorbent assay; GMT = geometric mean titer; ID_50_ = inhibitory dilution factor; IgG = immunoglobulin G; Nab = neutralizing antibody; ns, not significant; PBS = phosphate-buffered saline; PsVN = pseudovirus neutralization titer

High levels of binding antibody were elicited by vaccination in all animals 2 weeks after the first and second injection, with 4.5 to 9.4-fold increased S-2P binding titers measured after the second dose (Fig. 2b). Slightly lower antibody levels were observed for mRNA-1273.351 compared with mRNA-1273, potentially due to the coating S-2P protein used in the ELISA being homologous to mRNA-1273. These results demonstrate that both mRNA-1273.351 and mRNA-1273.211 are immunogenic in mice. mRNA-1273 elicited higher neutralization titers against the D614G than B.1.351 pseudovirus (Fig. 2c,d), although the approximate 2-fold difference was smaller than previously measured with phase 1 clinical trial sera [26]. mRNA-1273.351 elicited higher neutralization titers against B.1.351 compared with the D614G pseudovirus (Fig. 2c,d), with an approximate 4-fold difference in measured titers. When mRNA-1273.211 was used to vaccinate mice, similar neutralization titers were elicited against both the D614G and B.1.351 pseudoviruses (Fig. 2c,d), with no significant difference in neutralization titers. Thus, vaccination of mice with mRNA-1273.351 elicited high levels of neutralizing antibody against the B.1.351 pseudovirus and comparably lower levels versus D614G, whereas the multivalent mRNA-1273.211 vaccine stimulated robust neutralization responses against both D614G and B.1.351 pseudoviruses.

Sera from mice collected 2 weeks after the second injection was also assessed against SARS-CoV-2 variants that emerged in Brazil (P.1) and California (B.1.427/B.1.429 or CAL.20C). As described in Table 1, some of the mutations in these variants were different from both the Wuhan-Hu-1 and B.1.351 lineages, although the RBD mutations (K417T/N, E484K, N501Y) are common to both the P. 1 and B.1.351 viruses. As observed in previous assessments of NHPs and clinical trial sera [26], mice vaccinated with mRNA-1273 showed an approximate 2-fold reduction in neutralizing antibody levels against both the CAL.20C and P.1 variants (Fig. 2e). These reductions in neutralization titers against CAL.20C and P 1 variants were more pronounced in mice vaccinated with mRNA-1273.351 (3.7-fold and 2.6-fold reductions in geometric mean titers [GMTs] for mRNA-1273.351 compared with mRNA-1273, respectively). However, the multivalent mRNA-1273.211 vaccine neutralized these variants similarly to mRNA-1273.

### 3.2 mRNA-1273.351 was an effective third (booster) dose in animals previously vaccinated with a primary vaccination series of mRNA-1273

To evaluate the ability of the mRNA-1273.351 to boost pre-existing immunity and increase neutralization against both the wild-type and the B.1.351 virus, BALB/c mice were immunized with 1 or 0.1 μg mRNA-1273 on day 1 and 22, and the level and durability of the antibody responses were evaluated over the course of 7 months. Sera collection occurred on day 212, and a third dose of 1 or 0.1 μg mRNA-1273.351 was administered on day 213 (Fig. 3a).

**Fig. 3.**
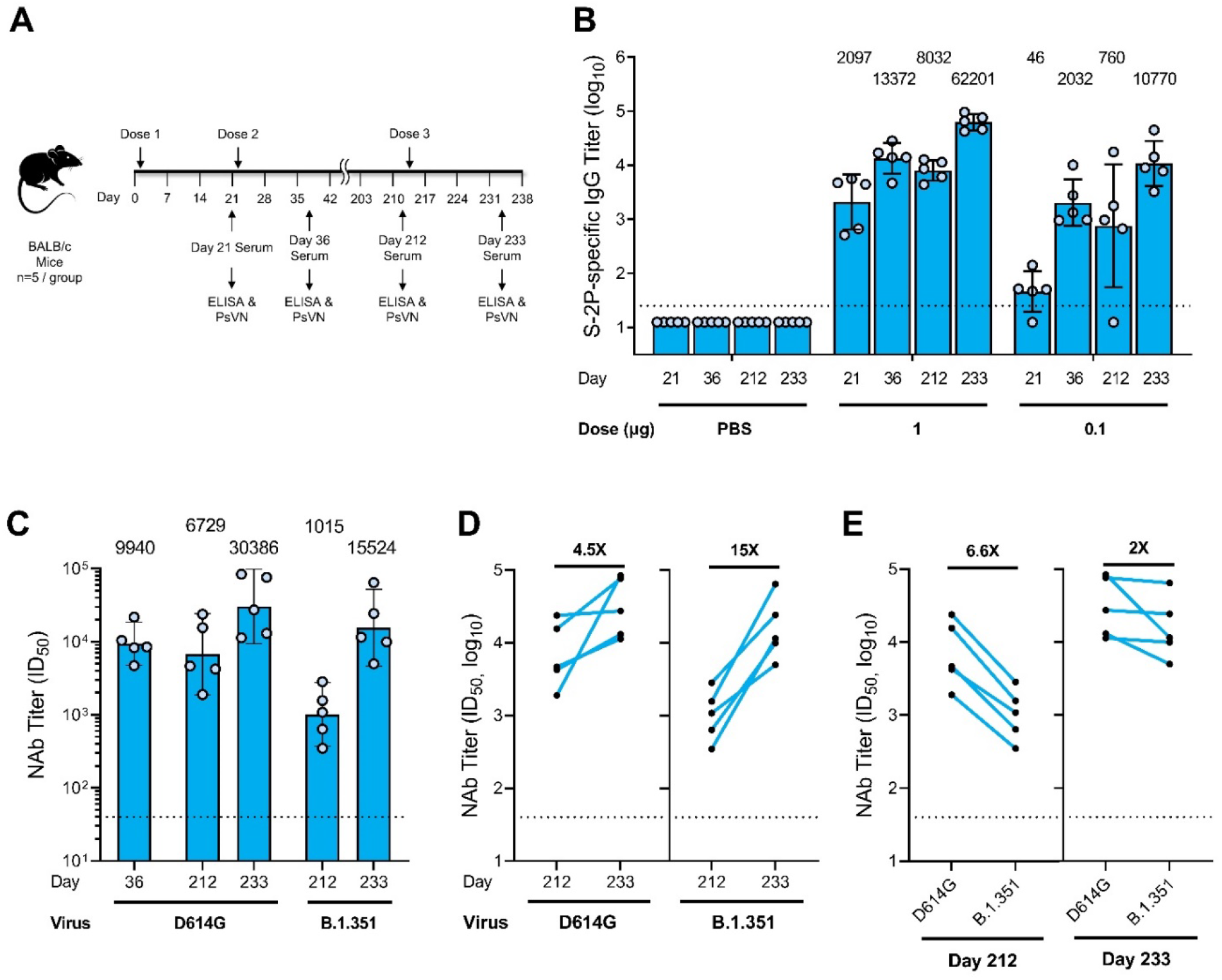
S-protein binding antibody and neutralization of D614G and B.1.351 SARS-CoV-2 pseudoviruses by serum from 1 μg mRNA-1273.351 boosted BALB/c mice. **a**, BALB/c mice were immunized with 1 or 0.1 μg mRNA-1273 (dose 1 on day 1; dose 2 on day 22) and were boosted with 1 or 0.1 μg mRNA-1273.351 on day 213. **b**, Results from individual mouse sera (n = 5 per group) are represented as dots on each figure, and the line is the mean of each group. The horizontal dotted line indicates the lower limit of quantitation for log_10_ IgG titer at 1.398. **c**, BALB/c mice previously immunized with mRNA-1273 were given a third dose with 1 μg mRNA-1273.351, with PsVN assessed against wild-type D614G and B.1.351 prior to dose 3 (Day 212) and 3 weeks after dose 3 (Day 233). Postdose 2 peak neutralization titer (Day 36) was referenced against D614G. **d**, Fold rise in neutralization against both viruses from the boosting dose of mRNA-1273.351. **e**, Fold difference in neutralization prior to and after dose 3. Postdose 2 peak neutralization titer reference (D614G assay). The box indicates the GMT, which is listed as text above each plot. The horizontal dotted line indicates the lower limit of quantitation for NAb titer at 40 ID_50_. Results from individual mouse sera is represented as dots on each figure, with lines connecting the D614G and B.1.351 neutralization titers. ELISA = enzyme-linked immunosorbent assay; IgG = immunoglobulin G; PBS = phosphate-buffered saline; PsVN = pseudovirus neutralization; GMT = geometric mean titer; ID_50_ = inhibitory dilution factor; NAb = neutralizing antibody.

High levels of binding antibody were elicited by vaccination with mRNA-1273, with peak titers measured 2 weeks after the second dose (Fig. 3b). After an initial drop in antibody levels, titers were stable over the 7-month monitoring period. Following the mRNA-1273.351 booster injection, antibody levels dramatically rose, exceeding the previously measured peak for both the 1 and 0.1 μg dose levels.

Neutralization titers were measured in the D614G PsVN assay on day 36 and 212, 1 day prior to the third dose. Titers remained high, with an ~1.5-fold drop measured over that period. Note that neutralizing titers at day 212 were measured in both the D614G and B.1.351 PsVN assays (Fig. 3c-e), with 6.6-fold higher titers measured in the D614G PsVN assay; this difference is similar to neutralization reductions observed with sera of NHPs and humans who received 2 doses of mRNA-1273 and increases the potential correlation of this animal model to what may be observed in humans.

The third dose of mRNA-1273.351 increased neutralization titers 4.5- and 15-fold in the D614G and B.1.351 PsVN assays, respectively (Fig. 3c,d). The difference between the 2 assays narrowed to 2-fold following the booster dose (Fig. 3c,e); the GMT of 15,524 against B.1.351 was ~1.5 fold higher than the peak titer against D614G 2 weeks after the second dose (Fig. 3c). Animals vaccinated at the 0.1 μg dose level had lower titers, but similar trends for binding antibody and neutralization titers were observed (data not shown).

## 4. Discussion

In this study, mRNA-1273.351 and mRNA-1273.211 were evaluated in mice as both a primary vaccination series and as a third booster dose in animals previously vaccinated with 2 injections of mRNA-1273. As a primary vaccination series, both vaccines were potently immunogenic after the first injection, with both S-2P binding and neutralizing antibody titers significantly increasing after the second injection. Neutralizing activity of mRNA-1273.351 against the D614G variant was 4-fold lower than that against the B.1.351 variant and 6.3-fold lower against D614G variant compared to mRNA-1273. In contrast, the multivalent mRNA-1273.211 vaccine elicited robust and comparable neutralizing titers against both D614G and B.1.351, which closely match those observed against D614G after mRNA-1273 vaccination. Thus, as a primary vaccination series, a multivalent approach appears most effective in broadening immune responses—as neutralization potency was enhanced against both B.1.351 and P.1 versus mRNA-1273 and remained significant against B.1.427/B.1.429/CAL.20C.

mRNA-1273.351 was evaluated as a booster injection in mice vaccinated with mRNA-1273 approximately 7 months previously. The third dose of mRNA-1273.351 dramatically boosted both S-2P binding antibody titers (Fig. 3a,b) and D614G and B.1.351 PsV neutralization titers (Fig. 3c-e). Neutralizing titers against B.1.351 PsV increased to levels well above the peak neutralizing titer against D614G after the second dose of mRNA-1273, the latter of which was fully protective in mice challenged with the mouse-adapted USA-WA1-F6/2020 variant [41]. In addition, the booster dose also increased neutralizing titers against D614G, although the fold-increase was less than that against B.1.351, as expected. Overall, the third injection of mRNA-1273.351 dramatically increased both D614G and B.1.351 neutralization titers, with titers much higher than the day 36 peak. Further, the difference in titers measured in the D614G and B.1.351 PsVN assays decreased from a 6.6-fold difference prior to the boosting dose, to a 2-fold difference 2 weeks after the third dose.

The number of animals available for boosting in this study allowed evaluation of only 1 boosting scenario (ie, mRNA-1273.351). Ongoing studies will evaluate the ability of mRNA-1273, mRNA-1273.351, and mRNA-1273.211 to effectively boost immunity driven by a primary vaccination series of mRNA-1273. Studies will also evaluate mRNA-1273.351 and mRNA-1273.211 in additional primary vaccination and boosting studies in mice, golden Syrian hamsters, and rhesus macaques, with either Wuhan-Hu-1 or B.1.351 challenge planned. These studies are designed to assess the level of neutralization of sera derived from vaccinated animals against pseudoviruses with either the Wuhan-Hu-1 D614G or the B.1.351 S proteins and the level of protection provided against Wuhan-Hu-1 and B.1.351 challenge. Global surveillance for the emergence of additional SARS-CoV-2 VOCs and the neutralization of VOCs by mRNA-1273 vaccinee sera are also ongoing. If additional variants emerge that reduce the neutralization capacity of mRNA-1273 further, additional mRNA vaccine designs may be developed and evaluated. In this study, the multivalent mRNA-1273.211 vaccine has already demonstrated to be an immunogenic strategy against multiple variants, and ongoing preclinical and clinical studies will provide further evidence of the utility of using a booster dose of mRNA-1273.211. This approach also supports seasonal adjustment, allowing for changes in response to the evolution of the SARS-CoV-2 virus.

Several potential limitations to the current study should be highlighted. Sera from mRNA-1273 vaccinated NHP or human sera was shown to have 6-8 fold reduced neutralizing activity against the B.1.351 variant SARS-CoV-2 in several assessments, although the level of neutralization remained at levels that are predicted to be protective [26]. Results in this study, however, showed that after the primary series of mRNA-1273, only a 2-fold reduced neutralization against the B.1.351 virus was evident 2 weeks after the second dose. A 6.6-fold reduction was seen at day 212, more relevant to what has been measured from human sera. Further studies in animal species more predictive of responses in humans such as NHPs as well as clinical studies in humans are ongoing. Further, only mRNA-1273.351 was evaluated as a third dose. The ability of the multivalent mRNA-1273.211 vaccine to boost immunity against both the D614G and B.1.351 viruses has not yet been assessed in preclinical models, although a recent exploratory analysis among a subset of patients in a clinical trial showed a mRNA-1273.211 booster dose increased neutralization against D614G and several VOCs, including B.1.351 [42]. Finally, very little antibody waning was measured in the 7-month evaluation of antibody levels in mice prior to the delivery of the boosting dose. This level of durability may not be reflective of that measured in NHPs or humans [42, 43].

The emergence of SARS-CoV-2 variants and the ability of the virus to partially overcome natural or vaccine-induced immunity served as a call to action. Not only are continued vaccination efforts needed to prevent the emergence of future VOCs, but strategies are needed for new SARS-CoV-2 vaccine research and development that can enhance the level of protection against key VOCs should they arise. The mRNA vaccine platform approach against SARS-CoV-2 VOCs has now been demonstrated in mice to be effective at broadening neutralization across variants and to boost antibody levels when applied as a third dose, with mitigation of the significant reduction in neutralization seen against the B.1.351 lineage. The mRNA platform allows for rapid design of vaccine antigens that incorporate key mutations, allowing for rapid future development of alternative variant-matched vaccines should they be needed. The designs evaluated in this study demonstrated potent cross-variant neutralization as a primary series and the ability to evolve the immune response through boosting; additional VOC designs can be rapidly developed and deployed in the future if needed to address the evolving SARS-CoV-2 virus.

## Abbreviations

ACE2: angiotensin converting enzyme 2
ELISA: enzyme-linked immunosorbent assay
GMT: geometric mean titer
ID50: inhibitory dilution factor
IgG: immunoglobulin G
LNP: lipid nanoparticle
Nab: neutralizing antibody
NHPs: non-human primates
ns: not significant
NTD: N-terminal domain
PBS: phosphate-buffered saline
PsVN: pseudovirus neutralization titer
RBD: receptor binding domain
RLUs: relative luminescence units
S: spike
SARS-CoV-2: severe acute respiratory syndrome coronavirus 2
TMB: tetramethylbenzidine
UTR: untranslated region
VOC: variant of concern
VOI: variant of interest
VSV: vesicular stomatitis virus

## Acknowledgements

We thank Michael Brunner and Dr. Michael Whitt for kind support on recombinant VSV-based SARS-CoV-2 pseudovirus production. Medical writing and editorial assistance was provided by Jared Cochran, PhD, of MEDiSTRAVA in accordance with Good Publication Practice (GPP3) guidelines, funded by Moderna Inc, and under the direction of the authors.

## Contributors

Conceptualization, K.W., S.E., S.R., E.N., G. S.-J., A.C., D.K.E.; data collection, K.W., A.C., M.K., S.E., L.M., D.L., A.W., C.H., C.P., A.H., H.J., J.Q., N.N., S.O., A.M.; analysis/interpretation of data, K.W., A.C., M.K., A.W., C.H., C.P., D.K.E.; writing—original draft preparation, D.K.E.; reviewing and editing, all authors. All authors have read and agreed to the published version of the manuscript.

## Data Availability

The authors declare that the data supporting the findings of this study are available within this article.

